# Second-order elasticities for Ecology and Evolution: Unravelling nonlinear fitness responses to perturbations

**DOI:** 10.1101/2025.04.30.651456

**Authors:** M. Kajin, S. Tuljapurkar, W. Zuo, H. Jaggi, S. Gascoigne, R. Salguero-Gómez

## Abstract

In ecology and evolutionary biology, understanding the relationship between vital rates (*e.g.*, survival, development, reproduction) and population growth is essential to elucidate how life history strategies are shaped by natural selection. However, the established demographic methods to decipher the relationship between vital rates and population growth often analyse only the linear changes in population fitness as a result of changes in vital rates, thus simplifying the complexities of said relationships. To overcome the widespread linearity simplification, here we introduce the second-order elasticities of mean population fitness, the *S-elasticity*. The S-elasticity quantifies how changes in one or more vital rates can produce a second-order change in mean fitness. We provide a systematic mathematical framework behind the S-elasticity, revealing its ability to identify the convex and concave responses of mean fitness to perturbations of vital rates. Through structured population models, we demonstrate the distinct roles of linear and nonlinear mean fitness responses, and their combination, enabling to characterise local concavity/convexity of the mean population fitness function. We illustrate the application and the differences of S-elasticities and their biological meanings using matrix population models of the armadillo (*Dasupys novemcinctus*) and Pyne’s plum (*Astragallus bibullatus*). These two case studies showcase how the S-elasticity provides key insights into mean fitness responses to perturbations on demographic process and their correlations. We discuss the improvements that the S-elasticity provides for species management and our understanding of how natural populations cope with environmental change.

## Introduction

In ecological and evolutionary dynamics, the various ways organisms go about completing their life cycles have produced a vast variation of life history strategies (Stearns 1992, Roff 2002). Said strategies have evolved over evolutionary time via natural selection (Gadgil and Bossert 1970), with environmental pressures having shaped how organisms optimise their vital rates of survival, growth, and reproduction (Morris and Doak 2004, McDonald et al. 2017). As such, life history strategies represent the evolutionary responses of organisms to their respective environments (Hutchings 2021). However, these life history strategies are not static, and continue to be subject to evolution today (Chase 1999, Jaggi et al. 2024). For instance, studies on guppies (*Poecilia reticulata*) have demonstrated rapid evolutionary changes in generation time, a dominant life history trait (Gaillard et al. 1989, Salguero-Gómez et al. 2016, Bassar et al. 2016), in response to predation pressure (Reznick and Endler 1982). Current and projected increases in environmental stochasticity (Boyce et al. 2006, Alexander et al. 2006) present a challenge to life histories that may have evolved under other environmental regimes (Cayuela et al. 2015, Paniw et al. 2018, Gascoigne et al. 2025). Thus, it is important to examine the evolutionary trajectories that have shaped these strategies and mitigate the contemporary pressures that may perturb them from their optimal states.

Life history strategies are governed by how vital rate values change across life cycle stages (Ebert 1999, Caswell 2001, Roff 2002). The stages (*i.e.*, distinct phases that an individual passes through from birth to death) can be defined by age, size, and/or development (Ebert 1999). The ecological, evolutionary, and conservation biology consequences of changes in life histories have been widely studied by examining the effects of small changes in vital rates (Zuidema & Franco 2001) using structured population models like the matrix population models (MPMs, Caswell 1978, Caswell et al. 1984, de Kroon et al. 1986, Crouse et al. 1987, van Tienderen 1995). The effect of such small changes (or perturbations) on the overall growth rate of the Matrix Population Model (MPM) is often described by its elasticity (Caswell et al. 1984, de Kroon et al. 1986). The growth rate of a population determines the average fitness of individuals in the population (Caswell 2001). This mean population fitness metric (fitness, hereafter) is shaped by the multiple matrix elements contained in an MPM. In this context, the usual elasticity measures the linear proportional response of fitness to a small proportional change in those matrix elements (de Kroon et al. 1986, Ebert 1999, Caswell 2001, Franco and Silvertown 2004).

Determining which demographic process –matrix element or underlying vital rate– has the highest impact on population growth rate has kept conservation biologists busy for decades. This focus on the most impacting demographic process was motivated by the application of MPMs to optimise management and harvest (Olmsted and Alvarez-Buylla 1995), conservation strategies (Steen and Erikstadt 1996, Heppell 1998, Wisdom et al. 2000), and pest control (McEvoy and Coombs 1999). Such an interpretation of the meaning of elasticity, that is which demographic process has the highest relative impact on population growth rate, operates within an ecological context. However, it is important to note that elasticities are local estimates of the effects of small relative perturbations on demographic processes on population growth rate (de Kroon et al. 1986, Ebert 1999, Caswell 2001).

Because the long-term population growth rate is a good proxy for the average, population-level fitness (Caswell 2001), elasticities represent a cohesive connection between ecology and evolution (Takada and Shefferson 2018). This connection arises because elasticities are closely related to selection gradients (de Kroon et al. 1986; Lande 1982; van Tienderen 2000). Selection gradients (Lande 1982), quantitatively expressed as the derivative of the fitness function with respect to a trait, indicate how fitness varies with trait values. As such, elasticities can represent a measure of the strength of natural selection acting on each demographic process (Takada and Shefferson 2018). This dichotomy of their interpretation –the ecological and evolutionary meanings of elasticity – places elasticities in a crossroad: that of eco-evolutionary dynamics (Shefferson and Salguero-Gómez 2015).

There is yet another interpretation of elasticities. Because elasticities can be obtained as partial derivatives of the population growth rate (Ebert 1999, Caswell 2001), their value represents the rate of change of fitness for the value of a specific demographic process (Fig. 1), and they represent scaled sensitivities. For instance, if the first-order partial derivative of the population growth rate with respect to a demographic process a_ij_ is positive 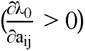, then fitness is expected to *increase linearly* as the mean value of a_ij_ across time increases, and *vice versa*: if 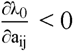 , then fitness is expected to *decrease linearly* as the a_ij_ mean increases. If 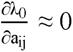, then changes in the mean of a_ij_ have no or little effect on fitness. Because elasticities are directly related to the first-order derivatives, we refer to elasticities as the *first-order effect on fitness*.

**Figure 1:**
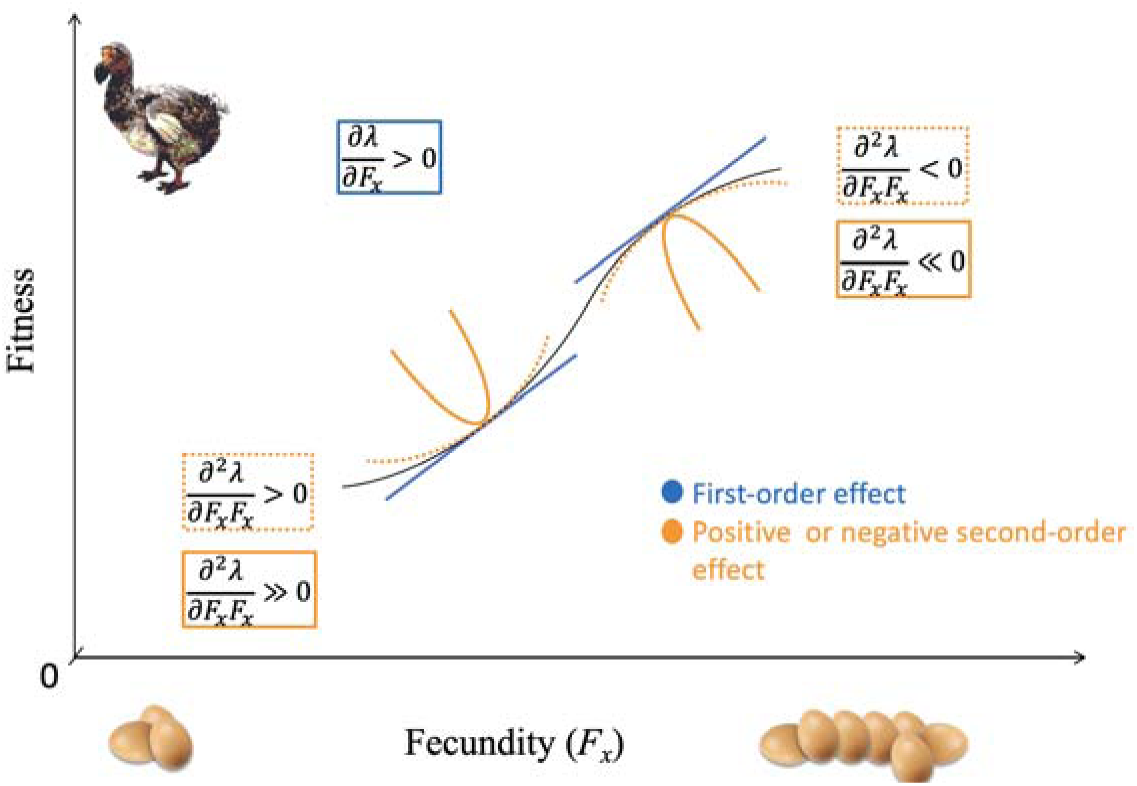
A theoretical representation of the fitness curve (black solid curve) associated with environmentally induced perturbations in single demographic process, namely fertility (F), for a hypothetical bird species with *x* age classes in the life cycle. The first-order effect of demographic process perturbations on fitness (blue solid line) represents the slope of the fitness tangent, indicating how a small perturbation in fertility results in a linear change in fitness, which corresponds to its elasticity. Fitness can only increase as *F_x_* increases, thus the slope of the fitness tangent can only be positive. The second-order effect represent the tangent’s curvature, capturing the nonlinear response of fitness to perturbations of fertility and directly relates to the S-elasticity we introduce in this manuscript. Negative second-order effects correspond to concave fitness shape, while positive second-order effects correspond to convex fitness shape. The intensity of the curvature is determined by the absolute value of the second-order effect (orange solid curve for strong, orange dashed curve for weak effect).

One can also describe a *second-order effect* of a small change in a demographic process of interest on fitness (Caswell 2001, Shyu and Caswell 2014). A second-order effect also measures the rate of change of fitness at a specific element similarly to a first-order effect. However, a second-order effect goes beyond the linearity assumption, revealing the *nonlinearity* of the fitness: whether fitness is *concave* or *convex* with respect to one or more demographic processes. Joining the information of a first-order effect on fitness with the information of a second-order effect permits identifying those conditions, where the second-order effect overrules the first-order effect. Importantly, the latter conditions help us determine where even small perturbations of demographic processes result in fast change in the population mean fitness (in both directions).

### Historical development of perturbation analysis and the resulting implications for evolutionary demography

When perturbation analysis methods started to be applied in population ecology, the focus was entirely on the first-order derivatives of population growth rate (Fig. 2). Hamilton (1966) famously examined the evolution of senescence in different demographic processes by obtaining the first derivatives of population growth rate (*λ*) with respect to each demographic process through the characteristic equation derived from the Euler-Lotka equation (Stearns 1992, Roff 2002), similar to Goodman (1971). Kato (1966) provided a rich background of the mathematical machinery behind perturbation analyses, which serves as a general guide in many scientific fields including population ecology (de Kroon et al. 1986, Ebert 1999, Caswell 2001). However, the term sensitivity was not used in population ecology until the late 1970s, when Caswell (1978) introduced the sensitivity of population growth rate to matrix elements (derived from vital rates) expressed in terms of left and right eigenvectors of the MPM (*i.e.*, reproductive value and stable stage distribution, respectively).

**Figure 2:**
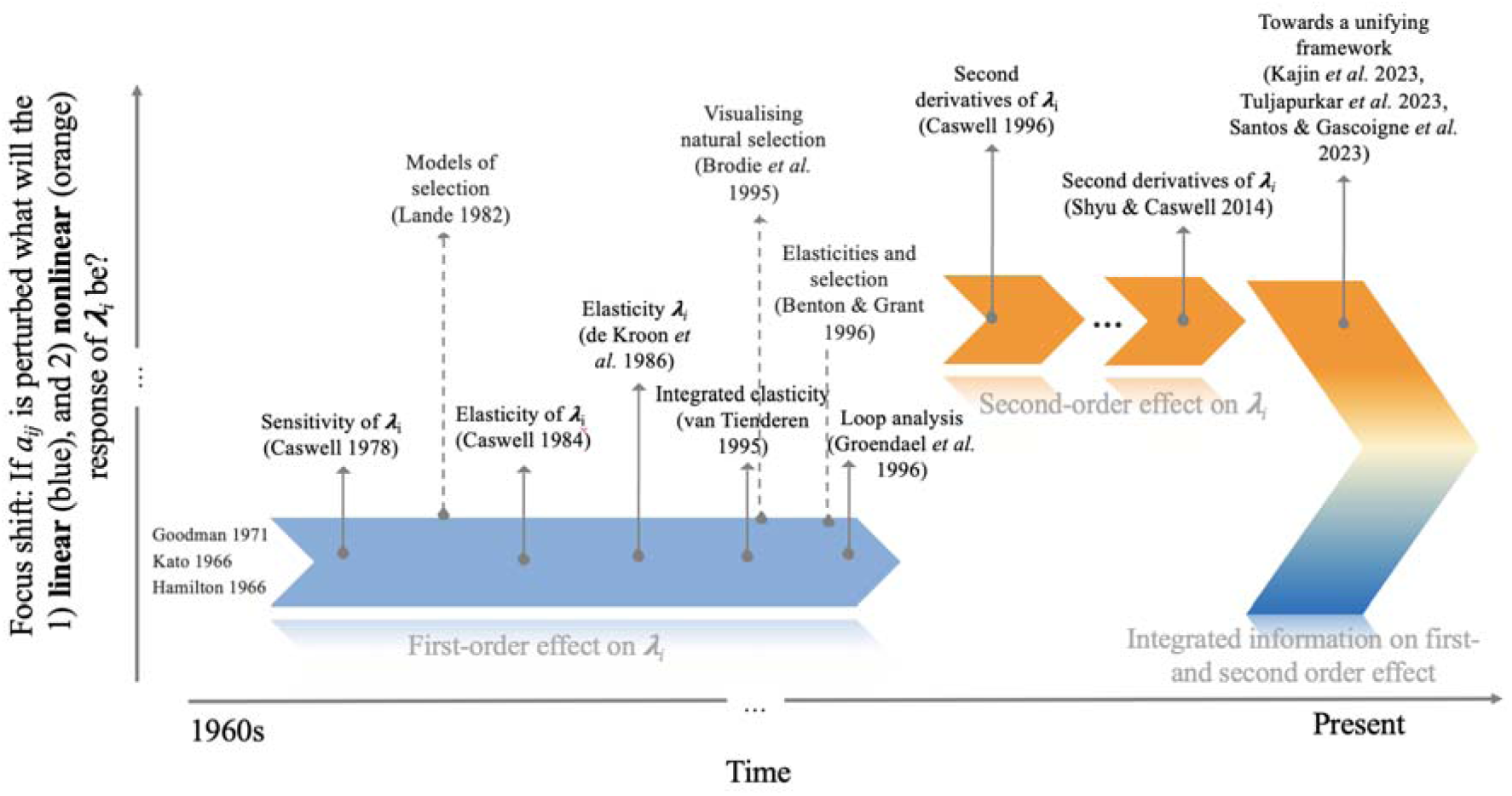
A non-comprehensive historic timeline of key publications on demographic perturbation analysis from 1960 to the present. The y-axis represents the layers of different types of information that a first- or second-order sensitivity analysis can offer. The blue arrow represents published papers that focused on a first-order effect of vital rate variability on fitness. The orange arrows represent the new layer of information added to the examination of the performance of populations: the second-order effect of variation in demographic processes on fitness. The three dots separating the red arrows (top-right) indicate the long period of time elapsed between the two main publications (Caswell 1996, Shyu and Caswell 2014) on the second derivatives of population growth rate (18 years). Studies that explicitly bridge ecological and evolutionary connotations of perturbation analyses in population ecology are indicated with dotted grey arrows, while ecologically oriented studies are designated by straight grey arrows. The blue-orange gradient represents the first attempts to report joint information on first and second-order effects of variation on population growth rate.

During the following two decades, several papers extended and improved the usage of sensitivities and elasticities, always focusing on a first-order effect of demographic process perturbation on fitness (Fig. 2). When Caswell (1996) published the second-order derivatives of population growth rate Click or tap here to enter text., the focus in population ecology widened from questions restricted by the linearity assumption to also ecological and evolutionary questions that could account for the nonlinear relationship between the fitness function and the demographic processes (Brooks et al. 2005, Carslake et al. 2008, Stott et al. 2012).

Reporting integrated results from linear and nonlinear fitness effect of fitness is both timely (Fig. 2) and necessary (Tuljapurkar et al. 2023). However, such integrated results are currently still under-developed in population ecology (Fig. 2; but see *e.g.* Santos et al. 2023, Tuljapurkar et al. 2023). The absence of an integrated linear and nonlinear approach in population ecology partly stems from the complexity of vector calculus involved in obtaining second order derivatives (Caswell 2001). The second-order sensitivity analysis has rarely been applied to plant and animal structured populations (but see Kajin et al. 2023; Shyu & Caswell 2014). Furthermore, the presence of -biologically inevitable-trade-offs (Roff 2002) between demographic processes has not yet received the attention it should, despite some important contributions towards the vital role that trade-offs play in sensitivity analyses and life-history theory (Charlesworth 1990, Jones and Tuljapurkar 2015).

To fill this gap, here we propose a definition of a novel second-order elasticity measure, which we coin as S-elasticity (where “S” stands for second order) and is denoted by 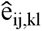. Our S-elasticity measures the second order fitness change resulting from a small proportional change in one or a pair of demographic processes a_ij_ and a_kl_ , where *i*, *j*, *k*, and *l* are the age, stage or size classes in a life cycle. We detail some key mathematical properties of the S-elasticity and show that this new measure has several key biological interpretations, including concavity/convexity of the fitness function. As such, the S-elasticity is an important complement to the toolbox of ecologists and evolutionary biologists examining the dynamics of population responses to increased environmental variation.

## Elasticity and S-elasticity

### Defining S-elasticity

In a discrete time and a constant environment setting, we can describe the dynamics of a structured population by a population matrix ***A***. Here, the population is structured into discrete stages that are represented by age classes, developmental stages, and/or discrete-size classes (Caswell 2001). The entries a_ij_ of the matrix ***A*** are matrix elements composed of vital rates (Zuidema and Franco 2001), the rates of transitions (from i to j) between ages, stages, and/or sizes (*e.g*., survival, development, *etc*.) and per-capita contributions (a/sexual reproduction). When the matrix ***A*** is primitive and irreducible, ***A*** has a dominant eigenvalue *λ_0_* with associated right *u*_0_ and a left eigenvector *v*_0_. In the context of elasticities/sensitivities, these eigenvectors are typically normalised so that their scalar product <*v*_0_,*u*_0_>= 1. When averaged across the population, the dominant eigenvalue of ***A*** represents the mean fitness of the population (*λ_0_*) described by ***A*** (Caswell 2001).

The behaviour of the mean fitness of a structured population can be studied using a matrix ***A***. Say that the environment or evolutionary pressures change a demographic process a_ij_ by a small proportion so that a_ij_ becomes a_ij_(1+ε). The elasticity of *λ_0_* to a_ij_ is the proportional *linear* change in *λ_0_* divided by the proportional change in a_ij_ (Caswell 1978, de Kroon et al. 1986). Because elasticities of matrix elements sum to unity across ***A*** (Ebert 1999, de Kroon et al. 2000, see also section Sum of S-elasticity below), they allow for a comparison of the relative contributions of each matrix element to the overall population growth rate across populations (Enright et al. 1995) and species (Franco and Silvertown 2004). However, the sum-to-unity rule implies that elasticities of *λ_0_* to matrix elements within MPM are not mathematically independent (Shea et al. 1994). When two matrix elements change, the sum-to-unity constraint means that change in a one matrix element is inherently driven by a change in the other matrix element (Van Tienderen 2000, Roff 2002, Doak et al. 2005, Jones and Tuljapurkar 2015). But this is a linear rule. When two matrix elements, say a_ij_ and a_kl_, change at the same time, the corresponding change in fitness is actually nonlinear. As a result, the change in fitness including this nonlinearity can be positive or negative depending on whether population growth is concave or convex. Thus, a comprehensive analysis of fitness behaviour should encompass both linear and nonlinear perspectives. The nonlinear perspective can be further divided into two cases:

1) To measure whether mean population fitness (*λ_0_*) is locally concave or convex for a *single* matrix element, we use the S-elasticity defined as

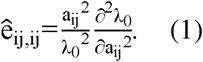

2) To measure whether mean population fitness (*λ_0_*) is locally concave or convex for a *pair* of matrix elements, a_ij_ and a_kl_ , we use the mixed S-elasticity defined as

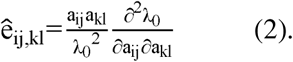

The nonlinear fitness response is determined by the sensitivity of two matrix elements and a response to a pulse disturbance (*i.e.,* sudden, one-time disturbance, Tuljapurkar et al. 2023). The response to said disturbance is determined by the second-order derivatives of population growth rate,

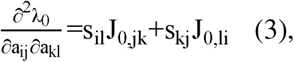

where the term s_ij_ corresponds to the sensitivity of population growth rate with respect to a_ij_

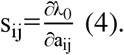

Because elasticities are proportional sensitivities (Caswell 1978), the corresponding elasticity e_ij_ of fitness (Ebert 1999, Caswell 2001, Tuljapurkar et al. 2023) is the proportional change that results in *λ_0_*,

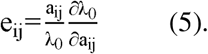

The ***J_0_*** matrix (Transient Response Matrix, TRM) (Eq. 3) quantifies the nonlinear selection pressures acting on demographic processes. Tuljapurkar et al. (2023) show how transient responses to pulse disturbances lead to a quantitative description of press disturbances and derive the connection from pulses to presses via TRM (***J_0_***). TRM can be used to understand how small, continuous changes (press disturbances *sensu* Menge and Sutherland [1976]) in demographic processes affect the long-term growth rate and structure of a population. TRM essentially captures the cumulative impact of these small changes over time, linking the immediate, short-term responses (transient dynamics) to the long-term, stable state of the population.

### Nonlinearity in Fitness

S-elasticities describe the nonlinear response in fitness to small, one-off pulse perturbations in demographic processes (Eq. 3). A change in the demographic process a_ij_ by a small proportion □_ij_ would cause mean population fitness to change from *λ*_0_ to *λ* such that:

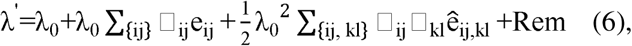

where the remainder *Rem* is of cubic order in the largest of the fractions □_ij_. Equation 6 can be examined by observing the linear and nonlinear terms in *λ* . The linear term is based on elasticities (e_ij_, Eq. 5), while the nonlinear term is based on the newly described S-elasticities for single elements (e_ij,ij_, Eq. 1) and for pairs of elements (e_ij,kl_, Eq. 2). To assess the relative impact of the nonlinear term compared to the linear term *for a pair of matrix elements* we can calculate the ratio of the S-elasticity to the elasticity as 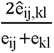. A high (positive or negative) S-elasticity/elasticity ratio indicates that the nonlinear term is stronger relative to the linear term in the fitness response when both a_ij_ and a_kl_ undergo small proportional changes. The sign of the S-elasticity/elasticity ratio for a pair of demographic processes –involved in a biologically possible trade-off– further indicates whether the mean population fitness is at peak of a concave surface (S-elasticity/elasticity ratio < 0), or at the bottom of a convex surface (S-elasticity/elasticity ratio > 0).

In a matrix population model ***A***, vital rate trade-offs (Roff 2002) are reflected by negative correlations between two lower-level matrix element parameters (Zuidema & Franco 2001). If two demographic processes in ***A***, say a_ij_ and a_kl_ , are involved in a trade-off, identifying the S-elasticities of a_ij_ and a_kl_ , becomes of interest. This is so because any perturbation in a_ij_ for example, will result in a_kl_ being changed simultaneously, thus the elasticities and the S-elasticities of a_kl_ will be influenced, even if a_kl_ itself was not perturbed directly. An informative way of identifying pairs of demographic processes involved in a vital rate trade-off is by obtaining the ratio between the elasticity of population growth rate to two demographic processes, 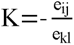 (Jones and Tuljapurkar 2015). The *K* ratio denotes the negative slope of the tangent line (thus the negative sign) on the iso-fitness curve, where the iso-fitness curve represents the different combinations of a_ij_ and a_kl_ that result in equal fitness (Jones and Tuljapurkar 2015).

Suppose mean population fitness is at an optimum before a change in two demographic processes. If fitness is at optimum, the nonlinear term of the change in fitness is negative. For the nonlinear term of the change in fitness to be negative, the following inequality must hold true for any feasible trade-off:

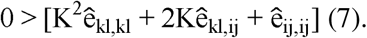

If the inequality holds, this means that the population is at a local peak in fitness. However, if the inequality does not hold, the population mean fitness might be far from its local optimum and the population might be going through a transient phase (Stott et al. 2011, 2012).

In this sense, we are looking for those pairs of demographic processes that are involved in a feasible trade-off (rather than discussing the shape of fitness curve for perturbations of single demographic process (as depicted in Fig. 1). The said pairs involved in a trade-off, where the second-order effect of perturbations on fitness (either positive or negative) overrules the first order effect on fitness, indicate that a small perturbation of either of the two demographic processes can lead to a fast change in fitness. On the other hand, near zero second-order effects on fitness identify those pairs where a substantial perturbation of either one or both demographic processes would be needed to cause a change in mean population fitness. Said differently, pairs of demographic processes with weak nonlinear terms (*i.e.,* near zero second-order effects) represent units of a life cycle where “variation can happen” – in the sense that variation in the said pairs of demographic processes will not cause harm to mean population fitness.

### Sum of S-elasticity

A second essential property of the S-elasticity lays in its summed effect. Elasticities are often grouped for biological and mathematical reasons to examine and classify impacts on life history strategies (*e.g.*, Franco & Silvertown 2004; Takada & Kawai 2020; Takada & Shefferson 2018). In many cases, life history transitions share a process (*e.g.,* growth), or an energy/mass flow (*e.g.,* reproduction), or describe a common outcome (*e.g.,* death). In such cases, elasticities are typically grouped together into vital rates (Grime 1977, Silvertown et al. 1992, Franco and Silvertown 2004, Takada and Kawai 2020).

Another way of grouping life history transitions defined by a matrix population model ***A*** is by following a life cycle loop (van Groenendael et al. 1994). Complex life cycles can be composed of many simple pathways (loops), where one loop may represent early reproduction and another loop late reproduction (Wardle 1998). By breaking down the life cycle of an organism into multiple constituent loops, we can identify life history strategies that most influence *λ*. Rather than focusing on a single demographic process, as classic elasticity analysis tends to do (e.g., Ebert 1999, Kroon et al. 2000, Caswell 2001), the loop approach provides a comprehensive understanding of the paths with the highest characteristic loop elasticities – which are particularly important for the long-term population growth rate.

Grouping S-elasticities also bears biological meaning. If every demographic process a_ij_ of the matrix ***A*** changes by the same small proportion □. Then, the new fitness becomes *λ* as per equation 6. Another way of thinking about the new fitness value following a change by the same proportion in all a_ij_ values means that the matrix population model changes from ***A*** to (1+ □)***A***. However, compared to the approach of changing individual demographic processes separately, the latter view simply means that mean population fitness changes from *λ*_0_ to:

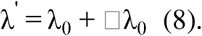

Because Equations 6 and 8 are identical, the following must hold:

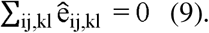

The S-elasticities summing to zero leads to the S-elasticities of different demographic processes not being independent of each other. Furthermore, if the Hessian (*i.e.*, the complete matrix of all the second-order derivatives) is not all 0 (trivial case), then the Hessian has at least one positive and one negative element (Sun and Sugie 2019). The latter yields the known result that the usual first-order elasticities of matrix elements sum to unity: ∑_ij_ e_ij_ =1 (de Kroon et al. 2000). The summative characteristics of elasticities and S-elasticities bear intriguing evolutionary implications, as they depict the distribution of importance in either ecological or evolutionary terms among various demographic processes in sustaining the long-term stability of population growth rates. The fact that the S-elasticities also sum to zero highlights their usefulness for comparative demography, across species and populations, and within populations across time and treatments.

### Example: Elasticities and S-elasticities for *Dasypus novemcinctus* and *Astragalus bibullatus*

To illustrate the insights that the S-elasticity can provide, we compare and contrast the elasticities and S-elasticities for a species of animal, the nine-banded armadillo (*Dasypus novemcinctus*, Mammalia, Dasypodidae) and a plant, the Pyne’s ground plum (*Astragalus bibullatus*, Angiosperms, Fabaceae). The demographic data for both species were obtained from the COMADRE (Salguero-Gómez et al. 2016) [24 Mar 2023, Version 4.23.3.1]) and COMPADRE data bases (Salguero-Gómez et al. 2015) [06 May 2023, Version 6.23.5.0]). We used the available MPMs in these databases, which consisted of three yearly MPMs for armadillo, and eight yearly matrices for the Pyne’s plum. The MPMs for both species are of 4×4 dimension and are based on size stages. We used the average matrices for both species, obtained from calculating the element-by-element arithmetic mean of the existing 3 MPMs for the armadillo and 8 MPMs for the Pyne’s plum. To show the joint dynamics of the elasticity and the S-elasticity, we calculated the population growth rate (*λ_0_*) for each MPM, elasticities (e_ij_), S-elasticities(ê*_ij,ij_* and ê*_ij,i_*ij,kl), the S-elasticity/elasticity ratios 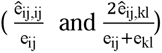 and the *K* ratio for pairs of matrix elements as described above using package *popbio* (Stubben et al. 2020) in RStudio (Posit team 2024).

## Results

The mean population fitness for both species is <1, namely λ_0_=0.868 for *D. novemcinctus* and λ_0_=0.940 for *A. bibullatus*, implying a projected decrease in population size by 13% and 6% annually, respectively, on the long-term, in the absence of density dependence and environmental stochasticity. In *D. novemcinctus*, the highest elasticity of *λ_0_* is to the 3^rd^ stage stasis (matrix element a ) (Fig. 3a), while in *A. bibullatus* the highest elasticiy of *λ_0_* is to 4^th^ stage stasis (a_4,4_) (Fig. 3b). Thus, mean population fitness is largely responsive to changes in adult stasis in both species.

**Figure 3:**
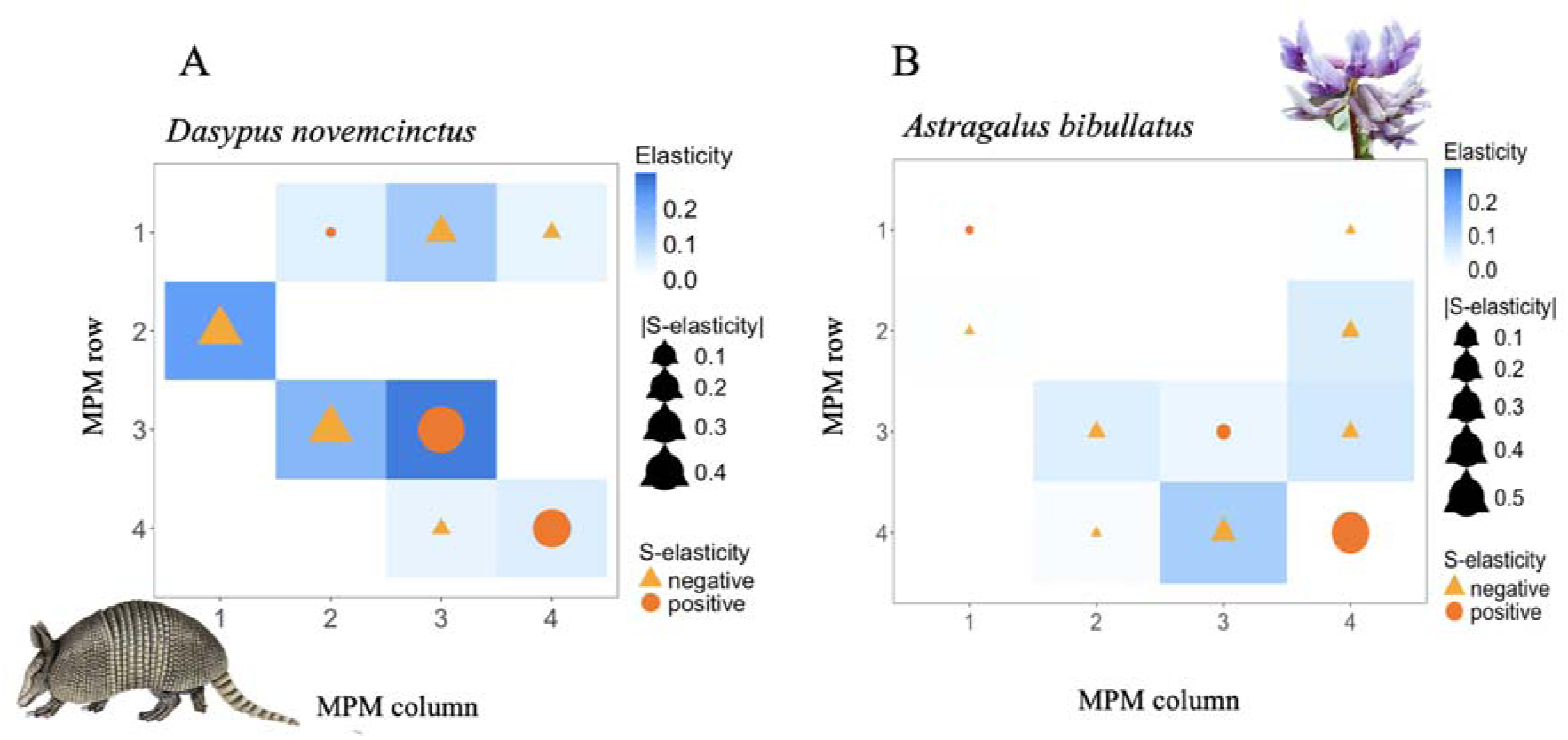
The role of demographic process perturbations on averaged mean fitness of individuals in a population as revealed by elasticities (first-) and S-elasticities (second-order effects of demographic process perturbation on fitness). First-order effects are represented by elasticities, while second-order effects are represented by the single-element S-elasticities of population growth rate (Eq. 1) for the averaged 4×4 matrix population models (MPM) of A) *Dasypus novemcinctus* and B) *Astragalus bibullatus*, from the COMADRE and COMPADRE databases, respectively. Each cell represents one matrix element where the *x* axis depicts the MPM column, and the *y* axis the MPM rows. The values of the elasticities of λ_0_ with respect to all matrix elements are represented by blue squared cells, while the values of the S-elasticities are illustrated by orange dots and triangles within the cells. Orange triangles denote negative S-elasticity, orange dots denote positive S-elasticity, and the size of the dots/triangles corresponds to the absolute value of the S-elasticity. White cells mean that the elasticity of λ_0_ to that MPM element is 0, while absence of a dot/triangle means that the S-elasticity is 0.

Displaying both first- and second-order effects on fitness alongside reveals some key similarities and differences between the two studied species. Indeed, Figure 3 illustrates that the demographic processes exhibiting the highest elasticities correspond to those demographic processes with highest |S-elasticity| values too. The nonlinear effects on fitness for *D. novemcinctus* and *A. bibullatus* are represented by the S-elasticities for a *single element* (Eq. 1, analogous to a self-second derivative) for each a_ij_ for each species (Fig. 3a and b, respectively).

Elements with highest elasticities display highest positive S-elasticities in both species. In *D. novemcinctus*, the matrix element a (3^rd^ stage stasis) results in highest elasticity and highest positive S-elasticity (Eq. 1, Fig. 3a). In *A. bibullatus*, the matrix element a (4^th^ stage stasis) results in highest elasticity and highest positive S-elasticity (Eq. 1, Fig. 3b). This means that fitness is locally convex for perturbations in element a_4,4_. Elements with second highest elasticities display negative S-elasticities. Namely, matrix elements a_2,1_ (growth from the 1^st^ to the 2^nd^ stage) and a (growth from the 2^nd^ to the 3^rd^ stage) in *D. novemcinctus* display high elasticity and negative S-elasticity (Eq. 1, Fig. 3a). This means that fitness is locally concave when only considering perturbations to a single demographic process.

The S-elasticities associated with pairs of demographic processes provide valuable insights into those processes that are involved in feasible trade-offs. Figure 4 joins the information of the pairs of demographic processes involved in feasible trade-offs with their respective S-elasticities. In *D. novemcinctus* the growth from the 3^rd^ stage to the 4^th^ stage (Fig. 4a, element a on the y-axis) and 4^th^ stage fertility (Fig. 4a, element a on the y-axis) are possibly involved in a trade-off with the 3^rd^ stage stasis (Fig. 4a, element a on the x-axis). Both pairs of elements (pair a_4,3_ and a_3,3_; and pair a_1,4_ and a_3,3_) display negative S-elasticities.

**Figure 4:**
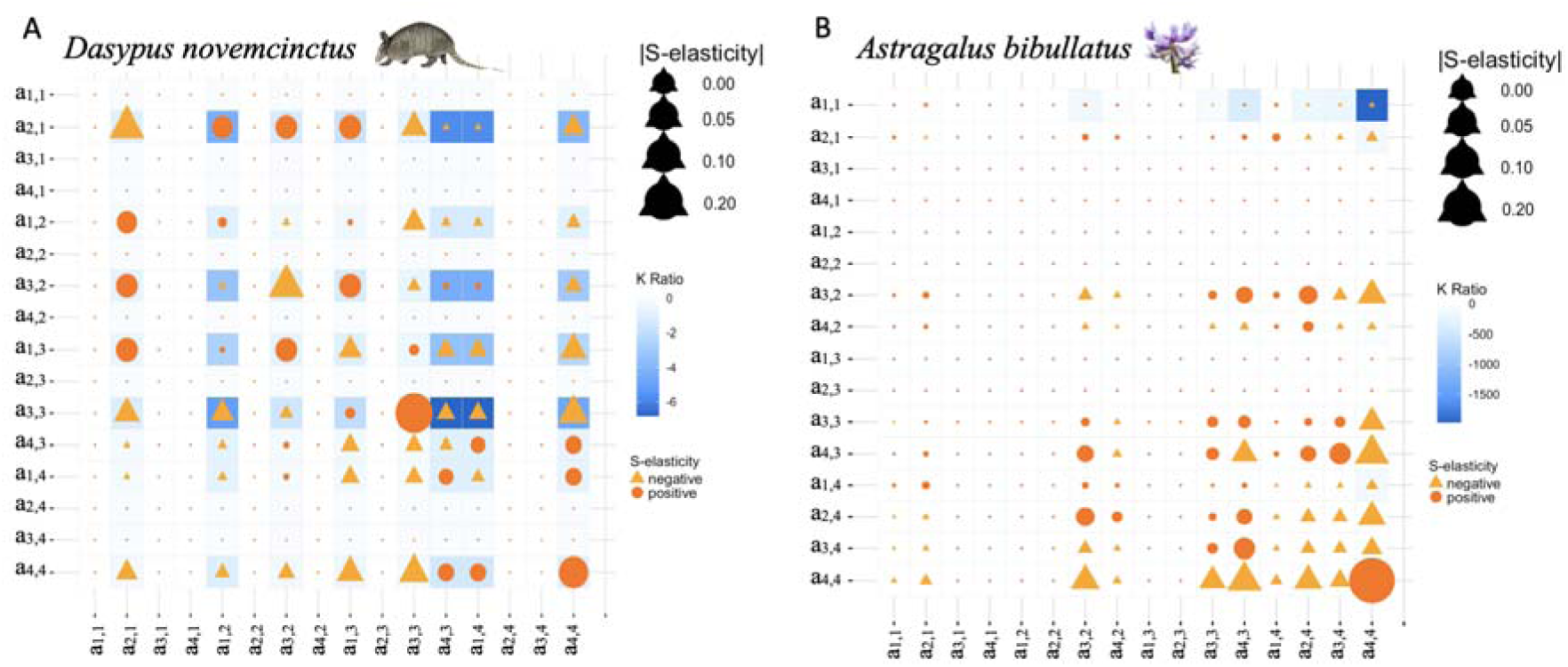
Second-order (*i.e.* nonlinear) effect of demographic process perturbation on fitness and feasible trade-offs between demographic processes. The *K* ratios and corresponding S-elasticities (Eq. 2) of fitness for a population of A) *Dasypus novemcinctus,* and B) *Astragallus bibullatus* for simultaneous changes in pairs of demographic processes. The 16×16 plot represents all pairwise combinations of matrix elements ordered by columns. The blue squares represent the *K* ratios for pairs of matrix elements, where the lower the ratio, the darker the colour. The orange dots/triangles represent the S-elasticities for changes in pairs of matrix elements: the orange triangles denote negative S-elasticities and the orange dots positive S-elasticities. The dot/triangle sizes are scaled by the absolute value of the S-elasticities, where the smaller the dot, the closer the value is to 0.

In *A. bibullatus*, the 4^th^ stage stasis is involved in a trade-off with 1^st^ stage stasis. This trade-off is identified by the value at the interface of element a_1,1_ on the y-axis and a_4,4_ on the x-axis (Fig. 4b). However, the near zero S-elasticity for the latter pair of demographic processes indicates that substantial perturbation of both demographic processes is needed to generate a change in mean population fitness.

We tested whether the inequality of Equation 7 holds for the pairs of matrix elements involved in a trade-off for both organisms. For *D. novemcinctus,* we investigated the inequality for the pair of elements a and a (*i.e.*, survival from the 3^rd^ to the the 4^th^ stage, and stasis in the 3^rd^ stage) and also for the pair a and a pair (fecundity of the 4^th^ stage and stasis of the 3^rd^ stage). For both pairs, the inequality holds as expected 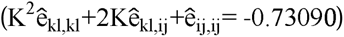. However, for *A. bibullatus*, the inequality for the a_j,1_ and a_4,4_ pair of matrix elements does not hold 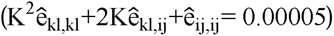.

The S-elasticity/elasticity ratio for pair of demographic processes 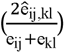 reveals where the second-order effect is strong compared to the first-order effect of perturbations in a_i,j_ and a_k,l_ on fitness. We plotted the S-elasticity/elasticity ratios in terms of their frequency to examine that, in *D. novemcinctus* (Fig. 5a), the second-order effect overrules the first-order effect (in the negative direction) since the highest absolute value of the S-elasticity/elasticity ratio is > 1 for the pair of matrix elements a_4,4_ and a_3,3_ (S-elasticity/elasticity ratio = -1.44). In contrast, for *A. bibullatus*, the second-order effect is weaker as compared to the first-order effect, since the highest absolute value of the S-elasticity/elasticity ratio is < 1 (Fig. 5b). Thus, in none of the matrix elements of *A. bibullatus* does the second-order effect of a perturbation on fitness overweight the first-order effect.

**Figure 5:**
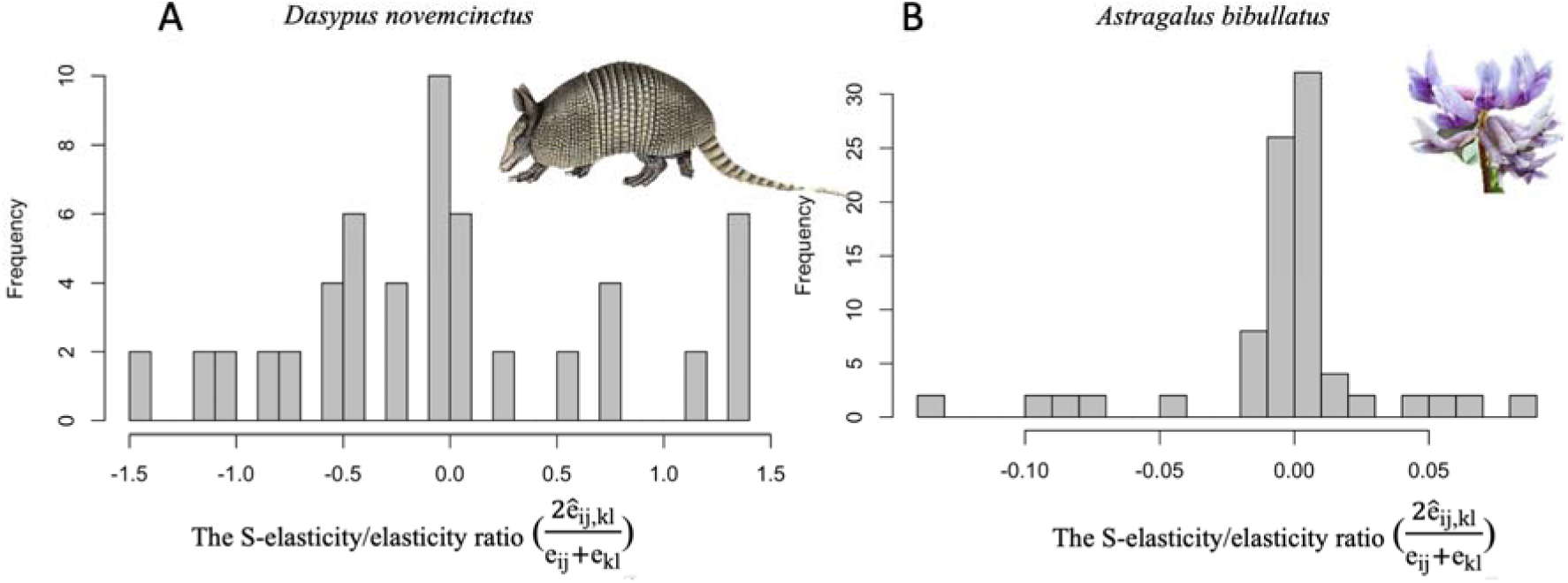
The strength of second- (*i.e.* nonlinear) relative to the strength of a first-order (*i.e.* linear) effect of demographic process perturbation on fitness. Shown are the frequencies of the S-elasticity/elasticity ratio for pairs of demographic processes in a population of A) *Dasypus novemcinctus,* and B) *Astragallus bibullatus*. Where the absolute value of the S-elasticity/elasticity ratio exceeds one, a second-order effect of perturbations on fitness overrules a first-order effect of demographic process perturbation on fitness.

## Discussion

We introduce a framework utilising S-elasticities to explore the nonlinear responses of mean population fitness to small perturbations in the elements of matrix population models. We show the relevance of studying higher-order effects of perturbations in matrix elements (or demographic processes) on population growth rate, and outline similarities and differences in S-elasticity patterns by using an animal (*Dasypus novemcinctus*) and a plant case study (*Astragallus bibullatus*, respectively). We discuss the implications of identifying local population-level fitness concavity and convexity for species management, while explicitly accounting for trade-offs between demographic processes.

The S-elasticities unlock a more integrative approach to study mean population fitness behaviour than linear approaches like elasticities (Ebert 1999, Kroon et al. 2000, Caswell 2001). The S-elasticities reveal how perturbations on demographic process may affect mean population fitness even when the assumption of linearity is relaxed. While other tools such as the integrated elasticities (van Tienderen 1995), for example, are a valuable upgrade towards an integrative approach, they still depend on the assumption of linearity between demographic processes and fitness. The S-elasticity does not only relax the linearity assumption but identifies the type of fitness nonlinearity: whether concave or convex. The introduced S-elasticity allows to identify those demographic processes where the strength of a second-order effect on fitness overrules the strength of a first-order effect of perturbations on mean population fitness. The said attribute is relevant for optimizing management strategies because it indicates for which pairs of demographic processes(es) and/or life cycle stage(s) the value of elasticity might be trivial. This triviality of elasticity may be due to the prevalence of strong non-linear fitness response over the linear response, either convex or concave. The importance of identifying nonlinear effects of perturbations on demographic processes on fitness has been long recognised (Carslake et al. 2008, 2009, Stott et al. 2012, Barraquand and Yoccoz 2013, Shyu and Caswell 2014, Tuljapurkar et al. 2023). The S-elasticity offers a straightforward tool to measure and compare the non-linear fitness responses inter- and intra-specifically.

The information that S-elasticity reveals about the 3^rd^ stage stasis for *D. novemcinctus* is two-layered. First, the 3^rd^ stage stasis is involved in trade off with late growth and late reproduction. Second, both latter pairs represent a local peak in fitness surface, meaning that perturbing these pairs of demographic processes would yield to a decline in population mean fitness. Depending on the situation, it might be logistically more realistic to prevent demographic process from being perturbed than attempting to increase their mean values (Ramula et al. 2008, Andrello et al. 2020). Therefore, by studying a higher- (*i.e*., second-) order effect of perturbations on fitness using S-elasticity, an alternative potential of an effective fitness optimum is revealed. Such a combined application of both first- and second-order effects of demographic process perturbation on fitness holds a potential for population ecology (Santos et al. 2023). This combined application indicates that elasticities can offer more valuable insights for species management, maximizing yield, or conservation efforts than simply identifying the demographic process with the highest importance for the population growth rate (Mills et al. 1999, Wisdom et al. 2000, Rojas-Sandoval and Meléndez-Ackerman 2013).

Examining the nonlinear mean fitness responses to perturbations in demographic processes can indicate a shift from concave to convex selection as environmental conditions change. Fitness shifts from concave to convex (or *vice versa*) are worth considering in the light of global climate change and habitat modification (Paniw et al. 2018, Albaladejo-Robles et al. 2022). This shift of selection type aligns with the broader eco-evolutionary framework, which proposes that rapid environmental changes necessitate equally dynamic and complex evolutionary responses (Gaillard et al. 2005, Shefferson and Salguero-Gómez 2015). The S- elasticities help pinpoint where fitness changes from concave to convex, or *vice versa*, for individual demographic processes (Eq. 1) or pairs of demographic processes (Eq. 2). By measuring local nonlinearity, S-elasticities can capture dynamic shifts on a fine eco-evolutionary scale.

Additional important feature of studying a second-order effect of perturbation on fitness is that it allows identifying the current stage of a studied population in terms of population stability (Farkas and Hagen 2007). The introduced inequality (Eq. 7) provides a diagnostic tool of whether a population is at a local peak in fitness or far from an optimum, a diagnosis that is of outmost importance when a strategy for management is being set up (Rozen et al. 2008, Jain et al. 2011, Gifford et al. 2011). The latter diagnosis combined with above detailed S-elasticity features provide a significant improvement to our understanding of how natural plant and animal populations might strategize and optimize their strive towards environmental unpredictability.

Future research should expand on the above findings by applying S-elasticity methodologies across various taxa and ecosystems to verify and expand upon these initial insights. Moreover, integrating long-term field studies and controlled experimental designs will provide empirical support for a better understanding of the behaviour of mean population fitness, validating how nonlinear selection and fitness responses emerge in natural settings (Doherty et al. 2004, Rodríguez-Caro et al. 2021). By introducing S-elasticities, we bridge significant gaps between classical ecological models and evolutionary dynamics. Our approach offers a refined perspective on ecological and evolutionary processes, facilitating a more nuanced understanding of population resilience and adaptability in an ever-changing world. Through such integrative strategies, we hope that ecologists and evolutionary biologists will be able to better anticipate and mitigate the impacts of rapid environmental change on biodiversity (Hamilton 1966, Tuljapurkar et al. 2023).

## Acknowledgements

MK was supported by the European Union via a Marie Curie Fellowship (MSCA MaxPersist #101032484) hosted by RSG, and NextGenerationEU # MN-0023-481. RSG was supported by a NERC Pushing the Frontiers grant (NE/X013766/1).

